# Direct segmentation of cortical cytoarchitectonic domains using ultra-high-resolution whole-brain diffusion MRI

**DOI:** 10.1101/2024.10.14.618245

**Authors:** Kristofor E. Pas, Kadharbatcha S. Saleem, Peter J. Basser, Alexandru V. Avram

## Abstract

We assess the potential of detecting cortical laminar patterns and areal borders by directly clustering voxel values of microstructural parameters derived from high-resolution mean apparent propagator (MAP) magnetic resonance imaging (MRI), as an alternative to conventional template-warping-based cortical parcellation methods. We acquired MAP-MRI data with 200*µ*m resolution in a fixed macaque monkey brain. To improve the sensitivity to cortical layers, we processed the data with a local anisotropic Gaussian filter determined voxel-wise by the plane tangent to the cortical surface. We directly clustered all cortical voxels using only the MAP-derived microstructural imaging biomarkers, with no information regarding their relative spatial location or dominant diffusion orientations. MAP-based 3D cytoarchitectonic segmentation revealed laminar patterns similar to those observed in the corresponding histological images. Moreover, transition regions between these laminar patterns agreed more accurately with histology than the borders between cortical areas estimated using conventional atlas/template-warping cortical parcellation. By cross-tabulating all cortical labels in the atlas– and MAP-based segmentations, we automatically matched the corresponding MAP-derived clusters (i.e., cytoarchitectonic domains) across the left and right hemispheres. Our results demonstrate that high-resolution MAP-MRI biomarkers can effectively delineate three-dimensional cortical cytoarchitectonic domains in single individuals. Their intrinsic tissue microstructural contrasts enable the construction of whole-brain mesoscopic cortical atlases.

## 1 Introduction

Partitioning the cortex into areas with distinct cyto– and myelo-architectonic features is fundamental to our ability to study the structure, function, organization, and development of healthy and diseased brains (Brodmann, 1909; Campbell, 1905; Vogt & Vogt, 1919; von Economo & Koskinas, 1925). This process, known as “cortical parcellation”, enables us to quantify disease-related morphological changes in specific cortical areas and to assess changes in the structure and function of large-scale brain networks. Moreover, cortical parcellation can provide a standard reference frame for neurosurgical navigation, focused ultrasound (FUS) delivery, radiation therapy planning, EEG electrode placement, and integration of multimodal brain activity studies.

Tissue domains with distinct cytoarchitectonic organizations, such as cortical layers and areas, can be studied in great detail using histological methods, such as histochemistry and immunohistochemistry. Because of their exquisite subcellular resolution and high specificity, these methods are considered the gold standard for delineating boundaries between cortical regions with distinct cytoarchitecture at the mesoscopic scale. However, histological techniques come with technical and logistical challenges (Amunts et al., 2013; Fischl & Sereno, 2018). They necessitate the analysis of tissue samples *ex vivo*, produce very large amounts of data, are time-consuming, expensive, and often have limited field-of-view (FOV). Nevertheless, despite these challenges, recent advances in data acquisition (Amunts et al., 2013; Shapson-Coe et al., 2024; Wagstyl et al., 2020) and computational processing (Paquola et al., 2019; Schiffer et al., 2021; Wagstyl et al., 2018, 2020) have greatly improved the throughput of histological processing and its feasibility for automatic analysis of large brain regions and cortical parcellation at the mesoscopic scale.

Current state-of-the-art methods for noninvasive, large-scale cortical parcellation rely on the diffeomorphic registration (or warping) of a subject’s brain MRI data to a standardized annotated template or atlas (Fischl, 2012). These methods are indispensable for analyzing data in cross-sectional and longitudinal whole-brain studies with large cohorts of human participants in both basic and clinical neuroscience research. The traditional approach involves warping a structural scan of the subject onto a standardized template with similar contrast, typically using *T*_1_-weighted or *T*_2_-weighted scans. The resulting deformation field is then used to transfer the cortical labels from the template to the subject’s brain. This process utilizes the 3D geometry of the cortical ribbon inferred from the structural MRI scans to drive cortical parcellation. However, since the conventional structural MRI scans lack sufficient contrast to cortical layers and areas, conventional atlas-based techniques do not incorporate subject-specific cytoarchitectonic features. As a result, while these methods can approximate boundaries aligned with prominent anatomical landmarks, such as the major sulci and gyri (e.g., central sulcus, Sylvian fissure, parieto-occipital sulcus), they are limited in their ability to parcellate the cortex into its finer specific areas and layers. When compared to histology (Amunts et al., 2007; Arslan et al., 2018; Fischl et al., 2008), template-warping-based cortical parcellation provides only approximate outlines of the subject’s cortical areal boundaries and may not accurately capture inter-individual variability (Amunts et al., 1999; Eickhoff et al., 2018). Given the essential function of cortical parcellation in neuroradiology, neurosurgery, neurodevelopment, and neuroscience, there is an urgent and critical need to develop more sensitive, accurate, and noninvasive methods for automated cortical parcellation based on endogenous contrasts having higher specificity to each subject’s intrinsic cortical cytoarchitectonic features.

Several studies have explored the possibility of optimizing various MRI contrast mechanisms, such as *T*_1_-weighted (Barazany & Assaf, 2012), *T*_2_-weighted (Glasser & van Essen, 2011), *T*_2_*-weighted (Kemper et al., 2018), MRI phase contrast (Duyn et al., 2007; Fischl & Wald, 2007), or magnetization transfer (Trampel et al., 2019) as a means to improve sensitivity to cortical cyto– and myeloarchitecture (Glasser & van Essen, 2011; Glasser et al., 2016), although results were mainly limited to selected cortical regions in the primary processing areas (Glasser & van Essen, 2011) and may require ultra-high field MRI scanners. Other studies have proposed parcellating the cortex using subject-specific information derived from large-scale structural or functional connectivity patterns (Barbas & Rempel-Clower, 1997) between brain regions that can be measured noninvasively using diffusion MRI fiber tractography (Anwander et al., 2007; Fan et al., 2016; Gao et al., 2018; Klein et al., 2007), or fMRI (Cohen et al., 2008; Craddock et al., 2012) respectively. Although these approaches showed promising results, they are not inherently sensitive to intrinsic cortical cytoarchitectonic features and may be prone to sources of bias in dMRI fiber tractography (Bastiani et al., 2013; Gao et al., 2018; Nie et al., 2011). Furthermore, discrepancies between cortical parcellations based on structural/functional connectivity and template warping remain unresolved (Amunts & Zilles, 2015; Zilles & Amunts, 2010).

Arguably, diffusion MRI (dMRI) provides one of the most sensitive and specific microstructural contrasts that can be acquired non-invasively. Diffusion MRI reflects important histological features, such as the size, shape, and density of cells (Pierpaoli & Basser, 1996), as well as myelin content (Budde & Annese, 2013) and the orientation of microstructural components (Basser et al., 1994a). Several studies using high-resolution diffusion tensor imaging (DTI) and high angular resolution diffusion imaging (HARDI) have shown good sensitivity to local areal boundaries and transition regions between laminar patterns observed with histology, especially for parameters characterizing diffusion anisotropy and descriptors of fiber orientation (Aggarwal et al., 2015; Assaf, 2019; Avram, Saleem, & Basser, 2022; Avram, Saleem, Komlosh, et al., 2022; Ganepola et al., 2018; Kleinnijenhuis et al., 2013; Leuze et al., 2014; McNab, Polimeni, et al., 2013; Nagy et al., 2013; N. Wang et al., 2020). Moreover, some studies suggest the possibility of using dMRI to quantify connectivity between cortical layers (Dell’Acqua et al., 2013; Leuze et al., 2014).

Mean Apparent Propagator (MAP) MRI (Özarslan et al., 2013) is an efficient, comprehensive, and clinically feasible (Avram et al., 2016) model-free method for quantifying water diffusion processes within tissues. In each voxel, MAP-MRI measures the probability density function of the net 3D displacements of water molecules diffusing in tissues, also known as diffusion propagators, generalizing and subsuming many other dMRI methods (Avram et al., 2017), such as DTI (Basser et al., 1994a; Basser et al., 1994b) and diffusion kurtosis imaging (DKI) (Jensen et al., 2005) while also providing a family of new microstructural parameters (Avram et al., 2016; Özarslan et al., 2013) that reflect local cytoarchitectonic features. Due to their excellent sensitivity to intrinsic cortical cytoarchitectonic features, high-resolution MAP-derived parameters have great potential for direct cytoarchitectonic segmentation and cortical parcellation.

In a previous paper, we investigated the spatial correlations between laminar patterns observed with high-resolution MAP-MRI-derived parameters and histology in a wide range of cortical areas (Avram, Saleem, Komlosh, et al., 2022). Despite differences in spatial resolution and biophysical contrast mechanisms between MAP and histological stains, we found similarities between laminar patterns measured with the two methods. However, we could not establish a one-to-one correspondence between the cortical layers observed with individual MAP parameters and histological stains that applied consistently across all cortical areas. To some extent, MAP parameters provide cytoarchitectonic information that can complement the characterization of cortical layers using conventional histological stains. Nonetheless, transition regions between MAP-derived laminar patterns seem to correlate well with the corresponding areal borders observed with histology (Avram, Saleem, Komlosh, et al., 2022).

In this study, we use an automated, data-driven clustering approach to classify cortical voxels directly based on cytoarchitectonic contrasts from multiple MAP-derived biomarkers simultaneously. We compare transition regions between MAP-derived cytoarchitectonic laminar patterns to those observed in histological stains and the warping-based cortical parcellation. Direct neuroimaging-based cytoarchitectonic mapping using endogenous microstructural contrasts could transform our ability to study the development, structure, function, and organization of the brain by improving accuracy and reducing bias resulting from inter-subject variability in cross-sectional and longitudinal studies with large cohorts of human participants.

## 2 Methods

### 2.1 Whole-brain high-resolution MAP-MRI of the macaque brain

We scanned the perfusion-fixed brain of a young adult macaque monkey (*Macaca mulatta*) using a Bruker 7T horizontal-bore MRI scanner and a Bruker 72 mm quadrature RF coil. We sealed the brain inside a custom 3D-printed plastic mold with a cylindrical enclosure (Saleem et al., 2021) and carefully aligned it inside the MRI scanner with the stereotaxic plane of the digital D99 macaque monkey brain atlas (Reveley et al., 2016), defined by the standard anatomical interaural axis and orbital ridges (Saleem & Logothetis, 2012). All procedures were performed under a protocol approved by the Institutional Animal Care and Use Committee of the National Institute of Mental Health (NIMH) and the National Institutes of Health (NIH) and adhered to the Guide for the Care and Use of Laboratory Animals (National Research Council).

We acquired MAP-MRI diffusion-weighted images (DWIs) at 200µm isotropic resolution using a 3D diffusion-weighted spin-echo (SE) echo-planar imaging (EPI) sequence with 50 ms echo time (TE) and 650 ms repetition time (TR). We used an imaging matrix of 375 *×* 320 *×* 230 on a 7.5 *×* 6.4 *×* 4.6 cm field-of-view (FOV) with 17 segments per k-z plane and 1.5 partial Fourier acceleration, and two averages for a scan duration of 2 hours per DWI volume. We acquired 112 DWIs using multiple b-value shells: 100, 1000, 2500, 4500, 7000, and 10,000 ms/mm^2^ with diffusion-encoding gradient orientations (3, 9, 15, 21, 28, and 36, respectively) uniformly sampling the unit sphere on each shell and across shells (Avram, Sarlls, et al., 2018; Koay et al., 2012). The diffusion gradient pulse duration and separation were *δ* = 6 ms and Δ= 28 ms, respectively. In the same session, we acquired magnetization transfer (MT) prepared MRI scans using a 3D gradient echo acquisition with 250 µm isotropic resolution (a 312 *×* 256 *×* 200 imaging matrix on a 7.8 *×* 6.4 *×* 5.0 cm FOV), a 15*^◦^* excitation flip angle, and TE/TR = 3.5/37 ms. The parameters of the MT saturation pulse were 2 kHz offset, 12.5 ms Gaussian pulse with 6.74 µT peak amplitude, 540*^◦^* flip angle. Two averages were obtained for each MT on and MT off scans.

### 2.2 Histological processing

After MR image acquisition, the brain was carefully blocked before histological work. By matching several anatomical landmarks on the surface of the brain with the corresponding locations on the 3D-rendered MRI volume we carefully adjusted the alignment of the brain to match the orientation with the coronal MRI sections, prior to sectioning, as described in our previous study (Saleem et al., 2021). We sectioned the entire brain into 50*µ*m thick coronal sections and prepared an alternating series of coronal tissue sections stained immunohistochemically with antibodies against neurofilament protein (SMI-32), parvalbumin (PV), and choline acetyltransferase (ChAT), or histochemically with Cresyl violet (CV) and Acetylcholinesterase (AchE) (Saleem et al., 2021). Finally, to compare cortical architectural features from histology and MRI, we obtained high-resolution images of all stained sections and individually matched each coronal histological section with a corresponding coronal slice from the MRI structural volume, as described in (Saleem et al., 2021).

### 2.3 MAP-MRI data processing and analysis

We computed the MT ratio (MTR) from images acquired with and without MT preparation to obtain a structural volume with excellent contrast between gray matter (GM) and white matter (WM). This MTR volume served as an anatomical template not only for preprocessing the diffusion MRI data but also for conventional warping-based cortical parcellation using the D99 digital macaque brain atlas (Reveley et al., 2016). Specifically, we processed all MAP-MRI data with the TORTOISE pipeline (Pierpaoli et al., 2010) using the MTR volume as the structural target for co-registration of all DWIs. The processing pipeline included Gibbs ringing artifact correction (Kellner et al., 2016), Marchenko-Pastur denoising (Veraart et al., 2016), image registration and correction of EPI distortions due to eddy currents, and *B*_0_ inhomogeneities.

To reduce noise and improve the visibility of the cortical layers, we designed an additional processing step for ultrahigh-resolution cortical dMRI data. Specifically, we denoised the corrected and co-registered MAP-DWI volumes using a local 3D anisotropic Gaussian filter (Perona & Malik, 1990) that smooths along the plane tangent to the cortical surface. For each cortical voxel, we estimated the orientation of the smoothing filter using only the MTR structural volume (Fig. 1). First, we segmented the cortical GM and brain WM using FSL-FAST (Y. Zhang et al., 2001). Next, using LayNii (L. R. Huber et al., 2021) we estimated the radial direction perpendicular to the cortical surface in each voxel, assuming equivolumetric layering (Bok, 1929; Waehnert et al., 2014). In each voxel, this radial direction implicitly determines its corresponding plane tangent to the orientation of the cortical surface, providing an approximate local cortical reference frame (CRF) within which both anisotropic diffusion signals (Avram, Saleem, & Basser, 2022; Avram et al., 2020a) and laminar signal profiles can be described efficiently (Waehnert et al., 2014).

**Figure 1:**
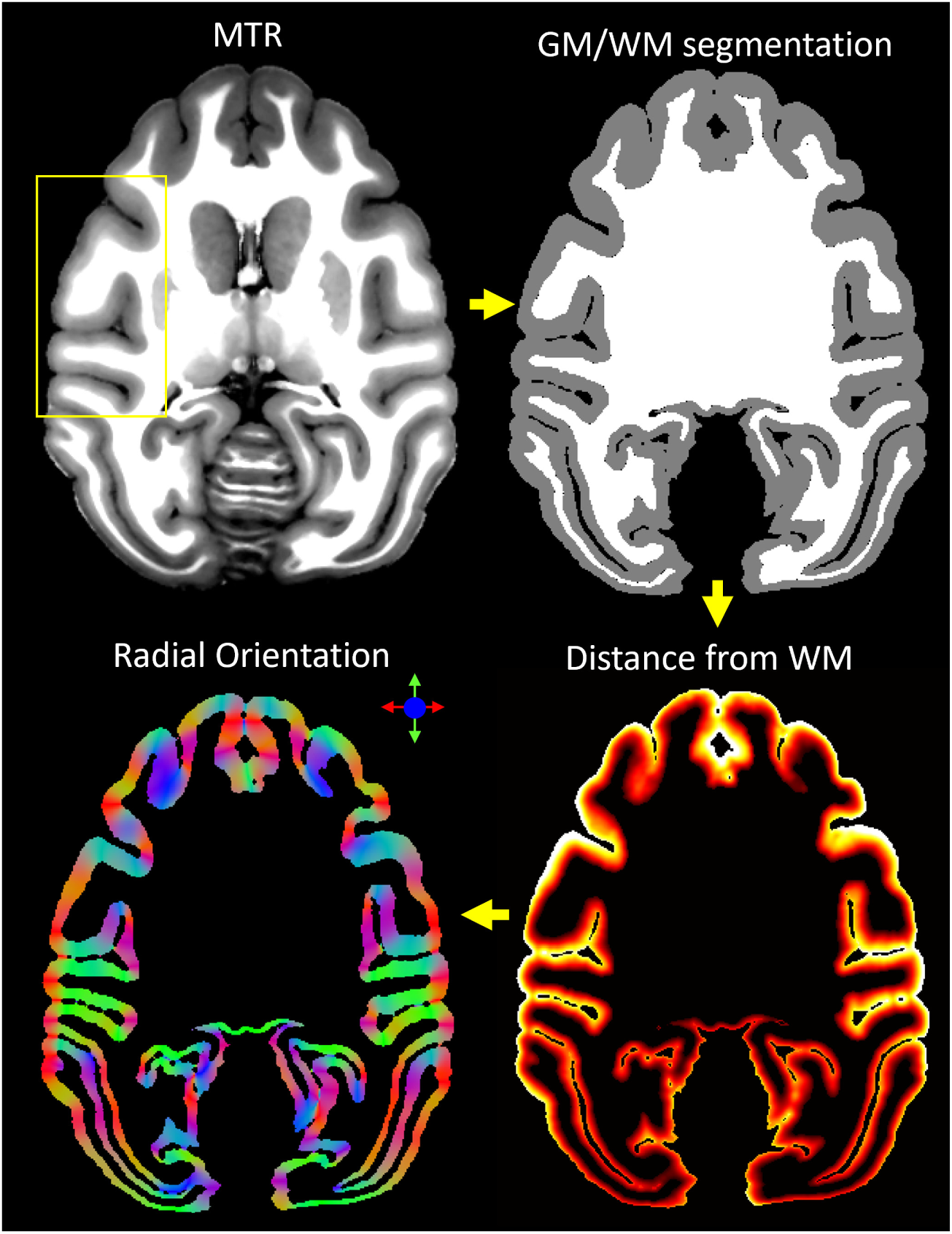
Local 3D anisotropic Gaussian filter-based denoising of cortical dMRI data using the cortical reference frame estimated from the structural MTR volume. From a structural magnetization transfer ratio (MTR) volume, we generated a white matter mask using FSL-FAST. Next, using LayNii, we computed the cortical depth of each voxel (i.e., distance from GM/WM boundary) and derived the direction perpendicular to the orientation of the cortical surface. This radial orientation of the cortical reference frame (CRF) implicitly defines the corresponding orthogonal plane locally tangent to the cortical surface, which determines the orientation of the local anisotropic Gaussian filter used to process the motion– and distortion-corrected MAP-DWIs.

We applied the local CRF-based anisotropic Gaussian denoising to each of the 112 distortion-corrected and co-registered DWIs separately. This step was implemented in MATLAB as a 3D convolution operation with a spatially varying 5×5×5 Gaussian kernel whose local orientation is determined by the CRF orientation in each voxel. The standard deviations were 100*µ*m and 1mm in the directions radial and tangential to the cortical surface, respectively. At each voxel location, the denoised value is computed as a weighted sum (3D Gaussian function) of the voxel intensities from the neighboring 5×5×5 voxels. Given the voxel size of 200*µm*, the maximum amount of blurring is 1mm in the plane tangent to the cortical surface (i.e., across areal borders).

From the CRF-filtered MAP-DWIs in the MTR space, we estimated the diffusion propagators by fitting the data in each voxel with a MAP-MRI Gauss-Hermite polynomial series expansion truncated at order 4. We performed the MAP-MRI analysis using the latest version of the TORTOISE software package (Irfanoglu et al., 2017) with default parameters as described in (Özarslan et al., 2013) Specifically, we enforced positive definiteness for the estimated propagator by sampling the displacement space on a 35×35×18 grid with the longest distance from the origin 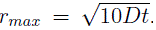. We used a Laplacian regularization constant of 0.02 (Fick et al., 2016). Subsequently, we computed scalar DTI-derived parameters, (i.e., fractional anisotropy – FA; mean, axial, and radial diffusivities-MD, AD, and RD, respectively) and MAP-MRI-derived microstructural parameters (i.e., propagator anisotropy – PA, non-Gaussianity – NG, return-to-origin probability – RTOP, return-to-axis probability – RTAP, and return-to-plane probability – RTPP).

### 2.4 Warping-based parcellation using the D99 macaque brain atlas

We performed conventional warping-based cortical parcellation using the same MTR volume that served as the structural template for DWI correction and co-registration. Specifically, we registered the MTR volume to the corresponding template of the D99 atlas using ANTs (Avants et al., 2009) accounting for both linear and multi-scale nonlinear transformations and used the resulting deformation field to warp the D99 atlas labels back to the MTR space.

### 2.5 Direct MAP-MRI-based cytoarchitectonic clustering

We clustered individual cortical voxels in each hemisphere (*≈* 2.3 million voxels per hemisphere) based on their values of scalar-valued MAP/DTI parameters (FA, RD, AD, PA, NG, RTOP, RTAP, and RTPP) using a Gaussian mixture model (GMM) clustering algorithm (McLachlan et al., 2019). The GMM algorithm provides a flexible probabilistic framework that handles large data sets effectively, captures complex data distributions, models uncertainty, is robust to outliers, and allows for cluster size variability. To prevent potential biases related to the geometry of the cortical ribbon (Bastiani et al., 2013; Nie et al., 2011), we did not include information about the relative voxel locations or underlying tissue architecture. Thus, the cytoarchitectonic clustering was restricted to the intrinsic microstructural features encoded in the scalar MAP parameters (e.g., anisotropy, non-gaussianity, zero-displacement probabilities) and was agnostic to any voxel information regarding the following:

1. The D99 cortical atlas labels estimated using warping-based parcellation
2. The relative spatial locations of individual voxels (i.e., spatial contiguity, or distance from white matter),
3. The underlying tissue orientation (i.e., the DEC map or fiber orientation distribution (FOD) glyphs).

The initial GMM clustering analysis was separately performed in each hemisphere using various combinations of MAP/DTI parameters as input features. For each choice of input features, the number of clusters was optimized using the Bayesian Information Criterion. The best segmentation consistency across the left and right hemispheres was obtained when the MAP parameters PA, NG, RTAP, and RTPP were used as features and the number of clusters was set to 14. Thus, the GMM clustering step produced a rough segmentation of cortical regions with 14 different MAP parameter signatures. To gain deeper insight into the relative utility of DTI-derived microstructural parameters we repeated the clustering for analysis using a set of input parameters that contained only FA, RD, and AD, and the same number of 14 clusters GMM clusters.

### 2.6 MAP-based cortical segmentation

Next, we used 3D morphological processing to separate disjoint (i.e., disconnected) components larger than 100 voxels initially assigned to the same GMM cluster and relabeled them as new clusters. The threshold of 100 contiguous voxels was chosen to comprise a net volume of 100 *×* 0.2 *×* 0.2 *×* 0.2*mm*^3^ = 0.8*nL*, roughly corresponding to the size of the smallest cortical cytoarchitectonic region that we expected to see. This represents the equivalent volume of a 3D cortical region with area 2*mm*^2^ and a layer thickness of 0.4*mm*, which is approximately the smallest cortical cytoarchitectonic domain observed with conventional histological methods (Brodmann, 1909). Isolated cluster components with volumes smaller than 100 voxels were likely due to noise and were merged with the neighboring cluster with the largest shared boundary. The 3D morphological processing step enables us to distinguish between cytoarchitectonic domains that have roughly the same MAP characteristic signature but are spatially disjoint. Moreover, this step reduced the dependence of the final segmentation on the number of clusters chosen in the initial GMM clustering analysis.

### 2.7 Matching MAP-based cortical labels in the left and right hemispheres

We automatically matched and labeled the cytoarchitectonic domains/regions segmented using GMM on the basis of voxelwise MAP parameter values in the left and right hemispheres. To achieve this, we first derived a canonical layer segmentation of each label in the symmetric D99 atlas using six layers with equivolumetric spacing obtained with LayNii (L. R. Huber et al., 2021) (Fig. 2). Then, for each hemisphere, we computed the cross-tabulation matrix between the MAP– and D99 layer segmentations and applied the Kuhn-Munkres assignment algorithm (Kuhn, 1955; Munkres, 1957) iteratively to match MAP labels of successively smaller size to the corresponding D99 layer label that showed the largest overlap. Cross-tabulation produces in each hemisphere a contingency matrix that counts the number of overlapping voxels between each pair of labels from two segmentations, similar to a two-dimensional histogram (Fig. 2). The rows of both contingency matrices correspond to the sorted D99 labels and are arranged in the same order, while their columns correspond to arbitrarily assigned MAP-labels. Permuting the rows and/or columns of the cross-tabulation matrix corresponds to a different ordering of the labels but characterizes the same overlap between the two segmentations. Because the D99 layer labels are symmetric, we can quantify the correspondence between the MAP labels in the two hemispheres by multiplying the transpose of the D99-MAP contingency matrix from the left hemisphere with the D99-MAP contingency matrix from the right hemisphere (Fig. 2). Processing and analysis steps for CRF-based filtering, GMM clustering analysis, 3D morphological filtering, and label matching were implemented in MATLAB 2023a.

**Figure 2:**
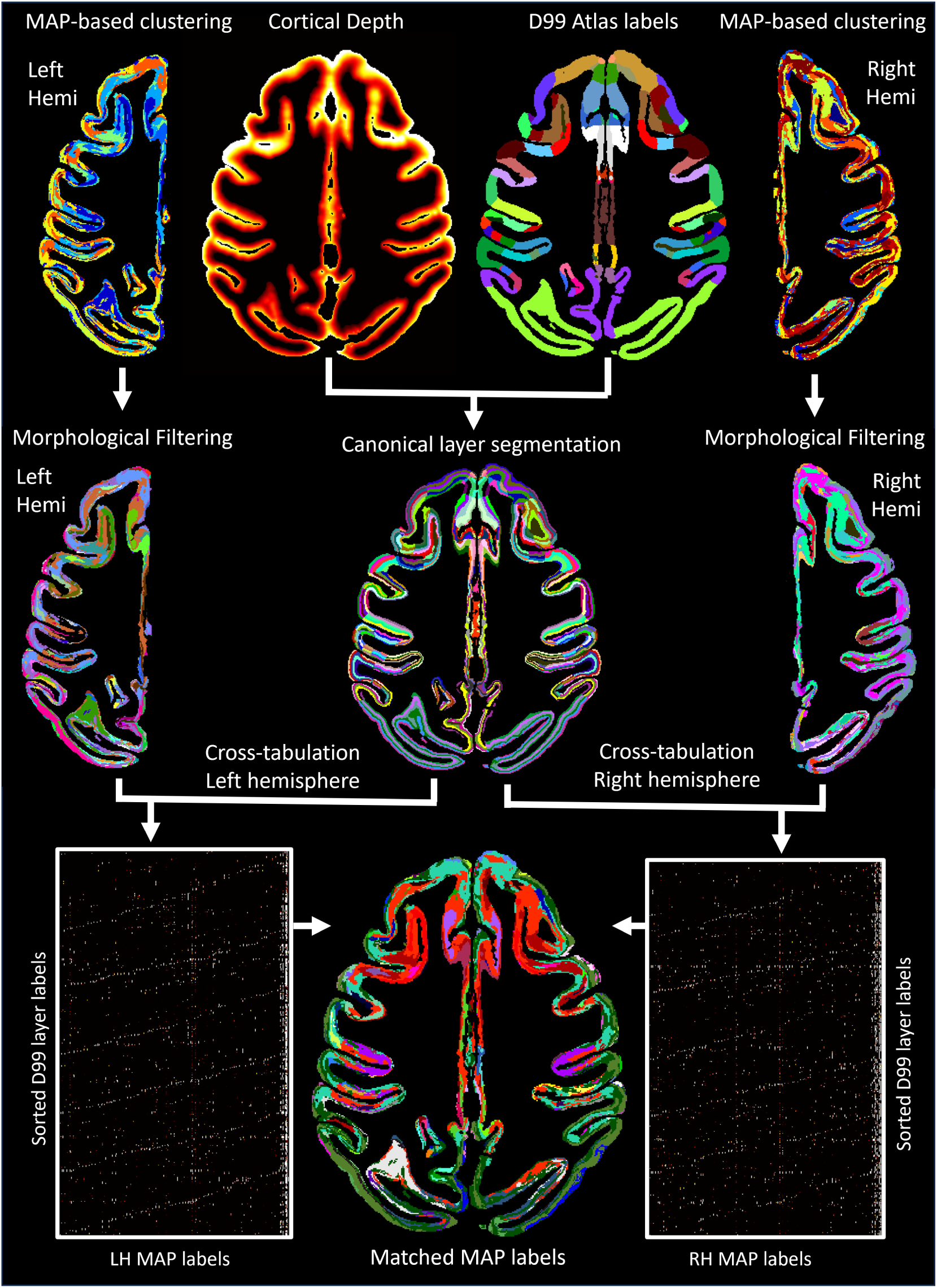
The outputs from MAP-based GMM clustering analysis in each hemisphere were processed using a 3D morphological filter, then cross-tabulated with the canonical layer segmentation derived from the symmetric D99 atlas in order to match the final MAP-based cytoarchitectonic labels in the left and right hemispheres.

### 2.8 Delineating boundaries between cortical areas

We evaluated the correspondence between the warping-based D99 cortical parcellation and direct MAP cytoarchitectonic segmentations and assessed their accuracy in identifying boundaries between cortical areas by comparing them with the corresponding histological images. Boundaries between cortical areas were manually delineated by an experienced neuroanatomist based on discontinuities in the laminar patterns observed in one or more histologically stained 2D coronal sections. During this process, the anatomist used the D99 atlas labels in standard space as an anatomical reference but was blinded to any subject-specific MRI-derived data including the MAP-MRI-derived parameter volumes and segmentations, as well as to the warped D99 atlas labels. The areal boundary demarcations were then superimposed on the matched coronal slice from the MTR volume to serve as a gold standard in the subsequent comparison with the MAP-based and warping-based segmentations. Since all dMRI data was co-registered to the structural MTR scan, the histologically defined areal boundaries can be directly compared to those derived using the MAP– and warping-based segmentations. In a few areas where the cortical geometry varied drastically through the coronal plane, the boundaries were carefully adjusted to account for slight misalignments in the cortical ribbon geometry across corresponding coronal sections from different histological stains and/or MRI slices.

## 3 Results

### 3.1 Cortical reference frame denoising of high-resolution dMRI data

MAP-DWI volumes showed an excellent signal-to-noise ratio (SNR) and negligible imaging distortions. They could be readily registered with the MTR volume. The outline of the segmented WM mask obtained from the MTR volume using FSL matched very well with the WM/GM boundary delineated by the high contrast of PA derived from the co-registered MAP-MRI data. Visual inspection and small manual edits/corrections of the WM and GM masks were performed prior to processing with LayNii (Fig. 1). The radial orientation that defines the CRF varied smoothly across the cortical ribbon. MAP and DTI parameter images computed from the CRF-filtered DWI volumes showed significantly less noise in cortical voxels and allowed a clearer visualization of laminar patterns (Fig. 3, green arrows) and discontinuities between them (Fig. 3, blue arrows).

**Figure 3:**
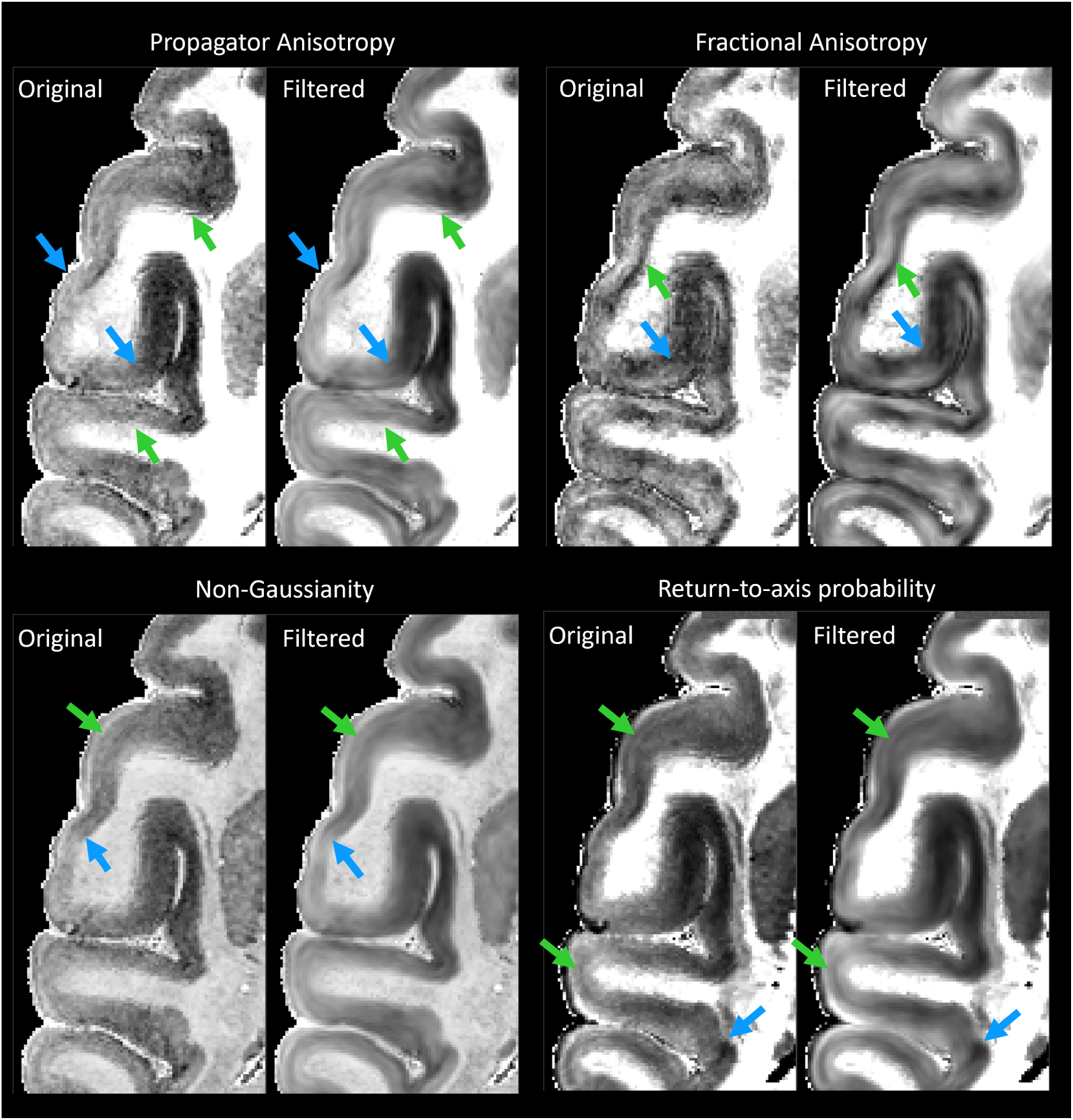
The effect of noise filtering using a local anisotropic Gaussian kernel on the estimation of MAP-MRI parameter volumes. MAP and DTI parameter images computed from the CRF-filtered DWI volumes showed significantly less noise in the cortical voxels and allowed a clearer visualization of laminar patterns (green arrows) and discontinuities between them (blue arrows).

### 3.2 High-resolution MAP-MRI of the cortex

Throughout most of the cortical gray matter, laminar patterns can be clearly observed on the high-resolution MAP-MRI parameter volumes (Fig. 5, green arrows). These laminar patterns correspond well with the cortical layers observed in the matched histological sections and have good symmetry between the left and right hemispheres (Fig. 5), which is in agreement with previous findings (Avram, Saleem, Komlosh, et al., 2022). The contrast of these laminar patterns appears different for various MAP parameters, further facilitating the discrimination of the underlying cytoarchitectonic boundaries. Transition regions between these laminar patterns can be observed in both the left and right hemispheres in good correspondence with the matched histological images (Fig. 5, blue arrows). Direct clustering of cortical cytoarchitectonic domains was performed using the values of these scalar-valued MAP parameters in individual voxels as features, without information about microstructural orientation, e.g., DEC-map.

**Figure 4:**
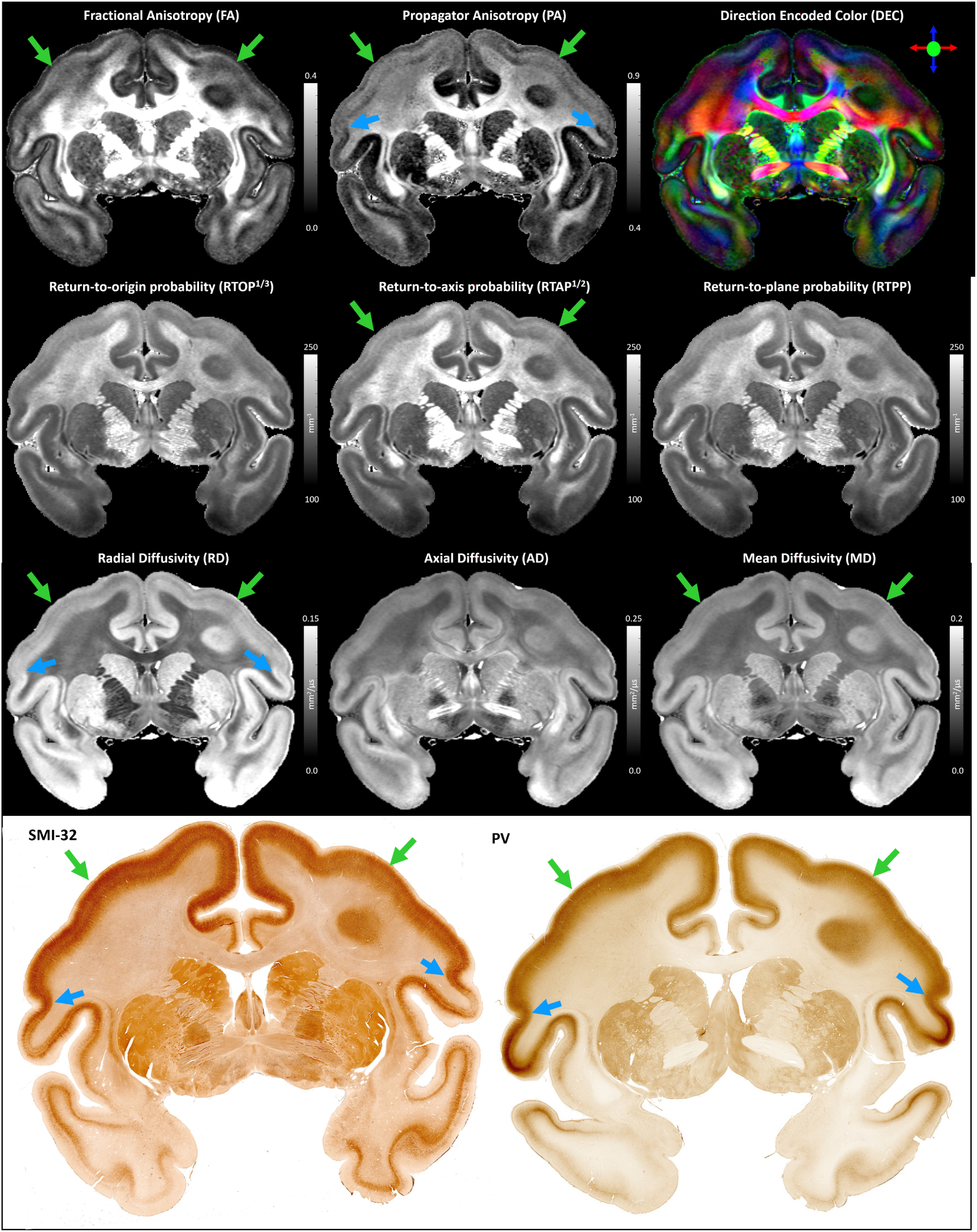
Coronal slices of the MAP-MRI parameters and matched histological images used in this study. The laminar patterns (green arrows) and discontinuities between them (blue arrows) are symmetric and in good agreement with histology. Direct clustering analysis using a Gaussian Mixture Model was performed in each hemisphere separately using the scalar-valued MAP parameters from individual voxels as features. The clustering analysis did not use information about the relative voxel locations or their microstructural orientations, e.g., vector-valued direction encoded color (DEC) map (top right).

**Figure 5:**
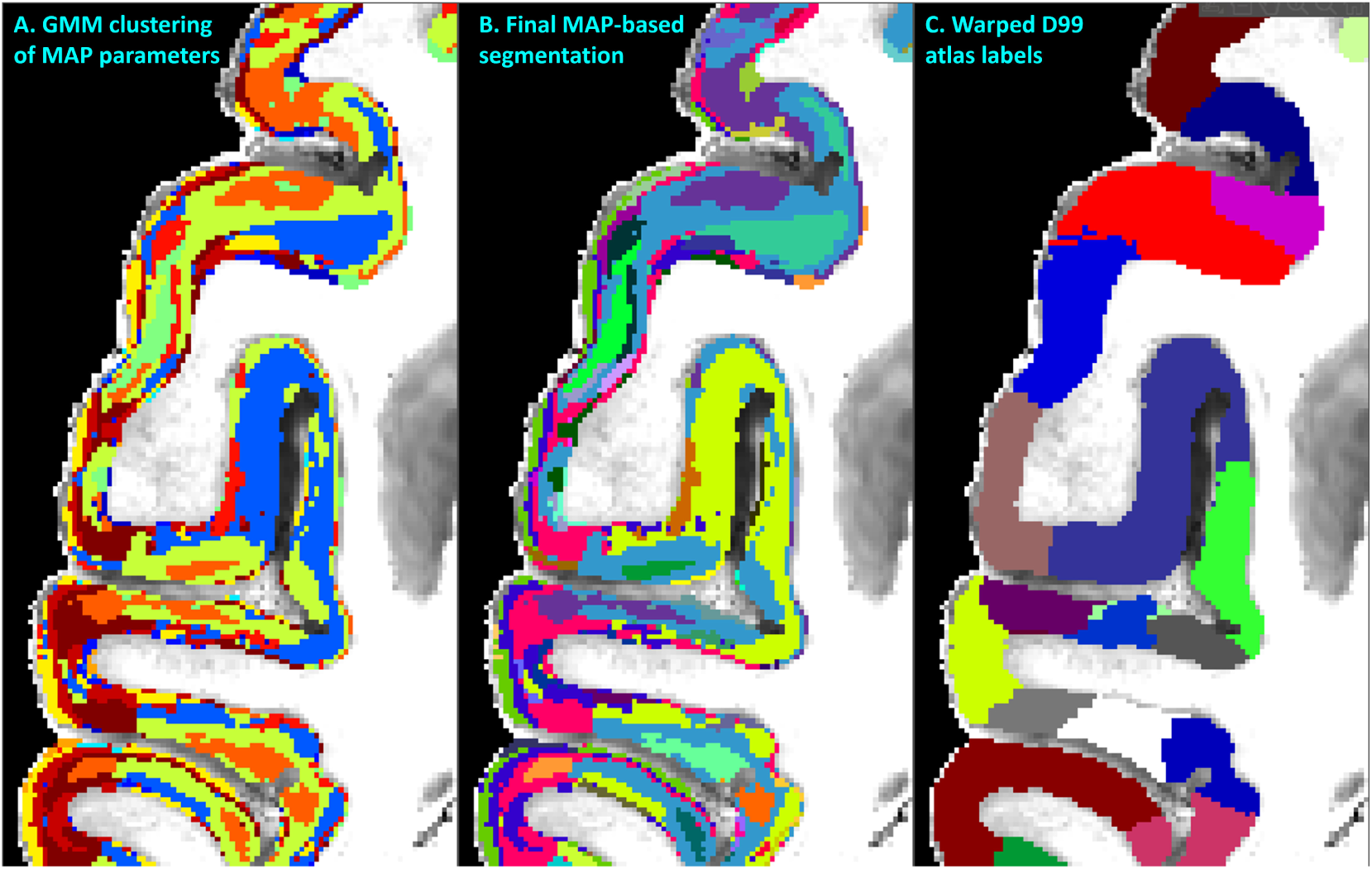
MAP-MRI parameter values from all cortical voxels in each hemisphere were segregated using GMM in 14 distinct clusters. (**A**). The resulting image was processed with a 3D morphological filter to merge small isolated spatial components/islands and uniquely relabel all spatially disjoint components, resulting in 420 distinct clusters per hemisphere. The final MAP-based segmentation **B** was obtained by matching labels between the left and right hemispheres. The final segmentation was compared with the corresponding warped D99 parcellation **C** and matched coronal histological sections. The same region from the axial slice in Fig. 3 is shown.

### 3.3 MAP-based direct 3D segmentation of cortical cytoarchitectonic domains

Despite the lack of spatial/orientational voxel information, direct voxel-wise GMM clustering prominently revealed spatially contiguous groups of voxels of different sizes and often with elongated shapes (laminar appearance) with thicknesses of *≈*200-800*µ*m (Fig. 5A). Such laminar patterns were observed when using various combinations of MAP and DTI parameters, although the best consistency/symmetry between the left and right hemispheres was obtained when PA, RTAP, NG, and RTPP were used as input features. The initial voxel-wise GMM analysis using these scalar MAP parameters yielded 14 distributed clusters with distinct cytoarchitectonic characteristics (Fig. 5A).

After the initial segmentation, isolated regions with volumes less than 100 voxels covered only 4.9% of the total cortical volume (i.e., the total number of cortical voxels). These small regions appeared isolated and localized at the periphery of the cortex, where tissue damage (tears/cuts) and air microbubbles were more likely to occur during sample preparation. Changing this 100-voxel threshold does not significantly impact the percent of voxels classified as artifactual/noisy that needed to be relabeled. For instance, using threshold values of 50, 200, and 500, the percentage of voxels categorized as noisy were 3.9% 6.2%, and 8.5% respectively, and did not significantly impact the final segmentation.

After 3D morphological filtering, the number of distinct clusters increased to *≈*420 in each hemisphere. The segmented clusters showed a high degree of spatial heterogeneity along the cortical ribbon of the subject with distinct laminar patterns and transition regions between them that generally correspond well with the canonical borders between cortical areas in the warped D99 parcellation (Fig. 5). Although these clusters were of different sizes and could not be uniquely correlated with specific histological stains, the MAP-based direct 3D cytoarchitectonic segmentation showed a high degree of symmetry between the left and right hemispheres. This symmetry could be observed more clearly after matching and labeling the MAP-based cytoarchitectonic domains in the two hemispheres to obtain the final segmentation shown in Fig. 6. Small incongruencies between the labels (i.e., colors) assigned automatically to these domains in the left and right hemispheres (Fig. 6) may be due to inaccuracies in the automatic matching process, given differences in the shapes and sizes of the corresponding regions across the hemispheres. Such anatomical differences may reflect important information regarding lateralization in cortical organization.

**Figure 6:**
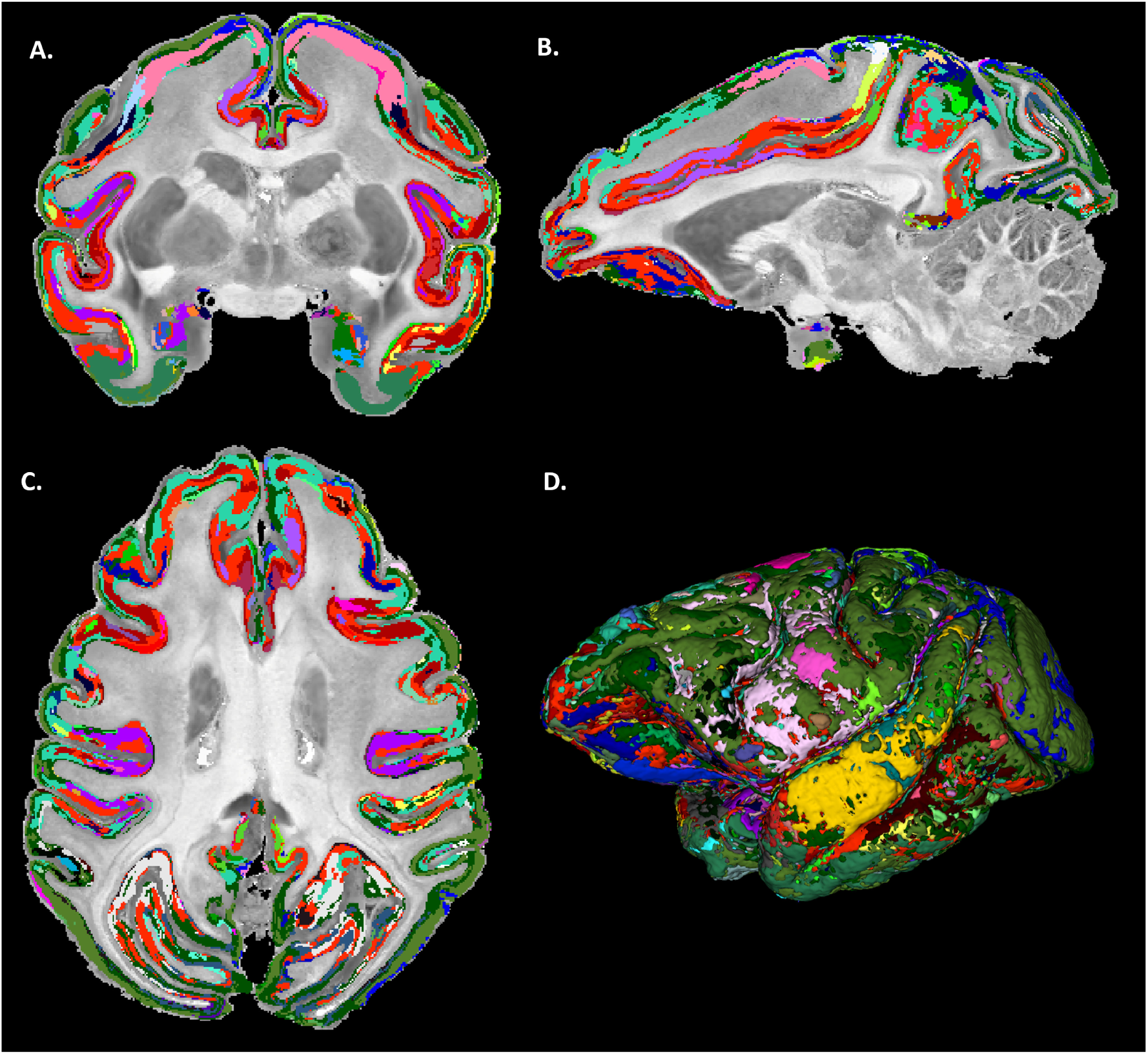
Final MAP-based cytoarchitectonic cortical domain segmentation with matched labels across the left and right hemispheres. A high degree of symmetry between the left and right hemispheres can be observed with MAP-based cytoarchitectonic domains roughly outlining the same specific layers and areas in both hemispheres (matched colors).

### 3.4 MAP-based cytoarchitectonic segmentation

The borders between adjacent MAP clusters are determined based on tissue microstructural and architectural differences as quantified in the normalized feature space of the MAP parameters. Thus, the borders between the cytoarchitectonic domains in the final MAP segmentation reflect transitions in the values of one or more MAP parameters. The MAP-based segmentation clearly revealed clusters at mesoscopic scale with laminar appearance, such as cortical layers, structures that are not present in the D99 atlas (Figs. 7-8). Some MAP cytoarchitectonic labels correspond to layers that extend/continue across areal boundaries often in good agreement with histology, whereas others terminate abruptly at the areal border. The sensitivity and specificity of individual MAP parameters to various cortical layers have been explored in a previous paper (Avram, Saleem, Komlosh, et al., 2022). MAP parameters quantify various features of water diffusion (modulated by the local tissue architecture), whereas histological stains directly measure the density of a particular molecule in the tissue. While there are no direct one-to-one correspondences between laminar patterns observed with MAP parameters and histological stains that apply universally across many cortical areas (Avram, Saleem, Komlosh, et al., 2022), MAP parameters such as PA, NG, RTAP, and RTPP provide a very informative characterization of cytoarchitectonic features across cortical layers, that can complement histological analysis. The PA quantifies how the mobility of tissue water varies along different orientations, whereas the NG provides a measure of tissue heterogeneity, including microscopic hindrances and compartments. Together, they are sensitive to general architectural and microstructural features of tissues. PA and NG typically show high values in the midcortical layers and, in certain cortical areas, provide laminar contrast that cannot be seen with traditional SMI-32 or PV staining (Avram, Saleem, Komlosh, et al., 2022). Meanwhile, zero-displacement probabilities, such as RTAP or RTPP, are sensitive to the average tissue water mobilities affected by microscopic hindrances and restrictions. They can correlate with laminar variations in cell densities and sizes of cells and can significantly improve cytoarchitectonic mapping in the superficial and deep layers (Avram, Saleem, Komlosh, et al., 2022).

**Figure 7:**
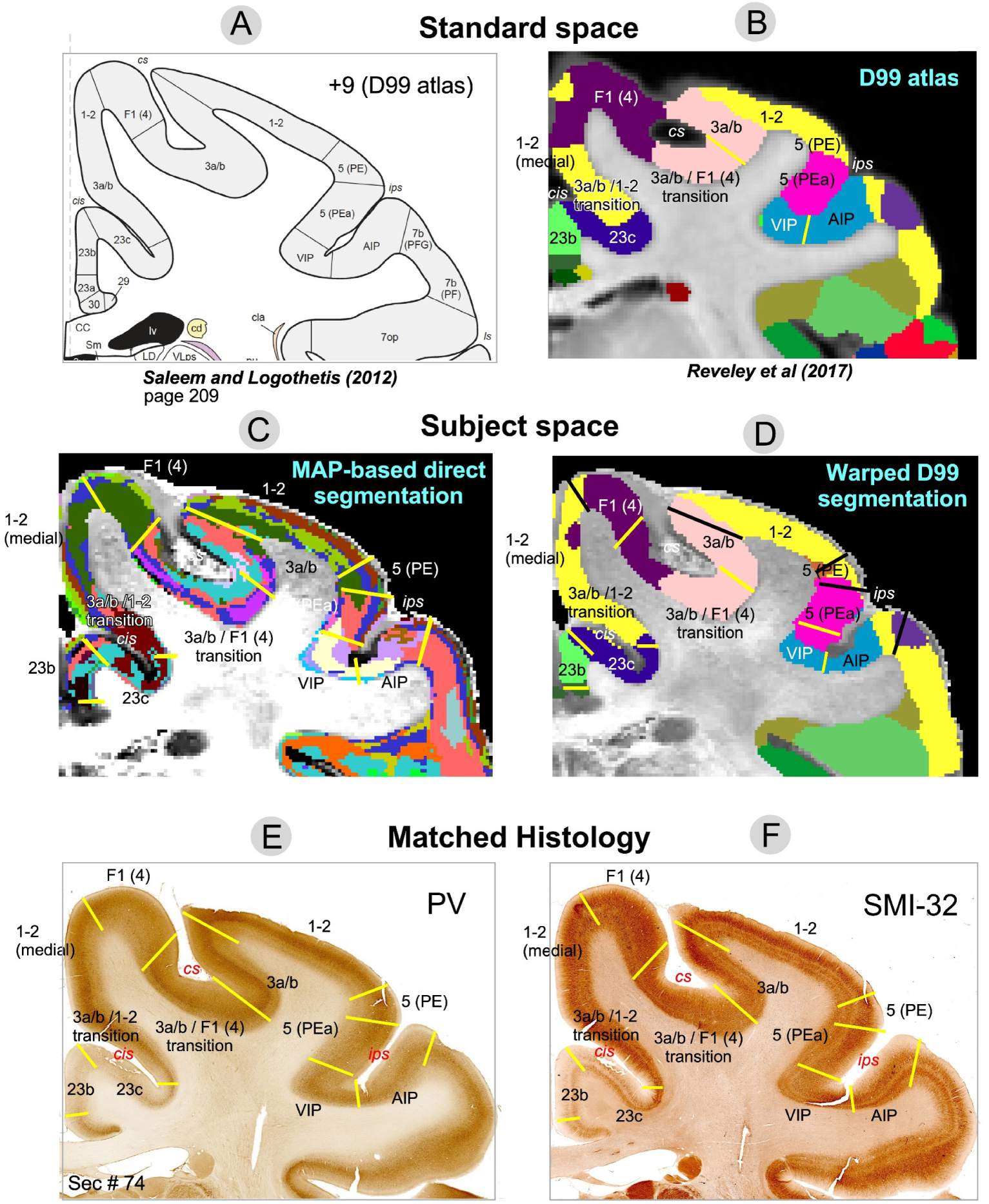
Comparison of warped D99 and direct MAP-based segmentations with two histological stains in a matched brain region. Boundaries between cortical areas were delineated manually based only on the histological images (E and F, yellow and black lines). Discontinuities in the laminar patterns in the MAP-based segmentation (C) correspond well with those observed in the PV (E) and SMI-32 (F) matched histological slices, providing a more accurate estimation of areal boundaries than the warping-based segmentation (D); e.g., the borders between area F1(4) and the 3a/b-F1(4) transition region, or between areas 1-2 and 5(PE), 5(PEa) and VIP, among others. The best correspondences between the warping-based D99 areal boundaries and the matched histology were generally found in regions of maximum cortical curvature, e.g., the boundary between areas 1-2 and 3a/b.

**Figure 8:**
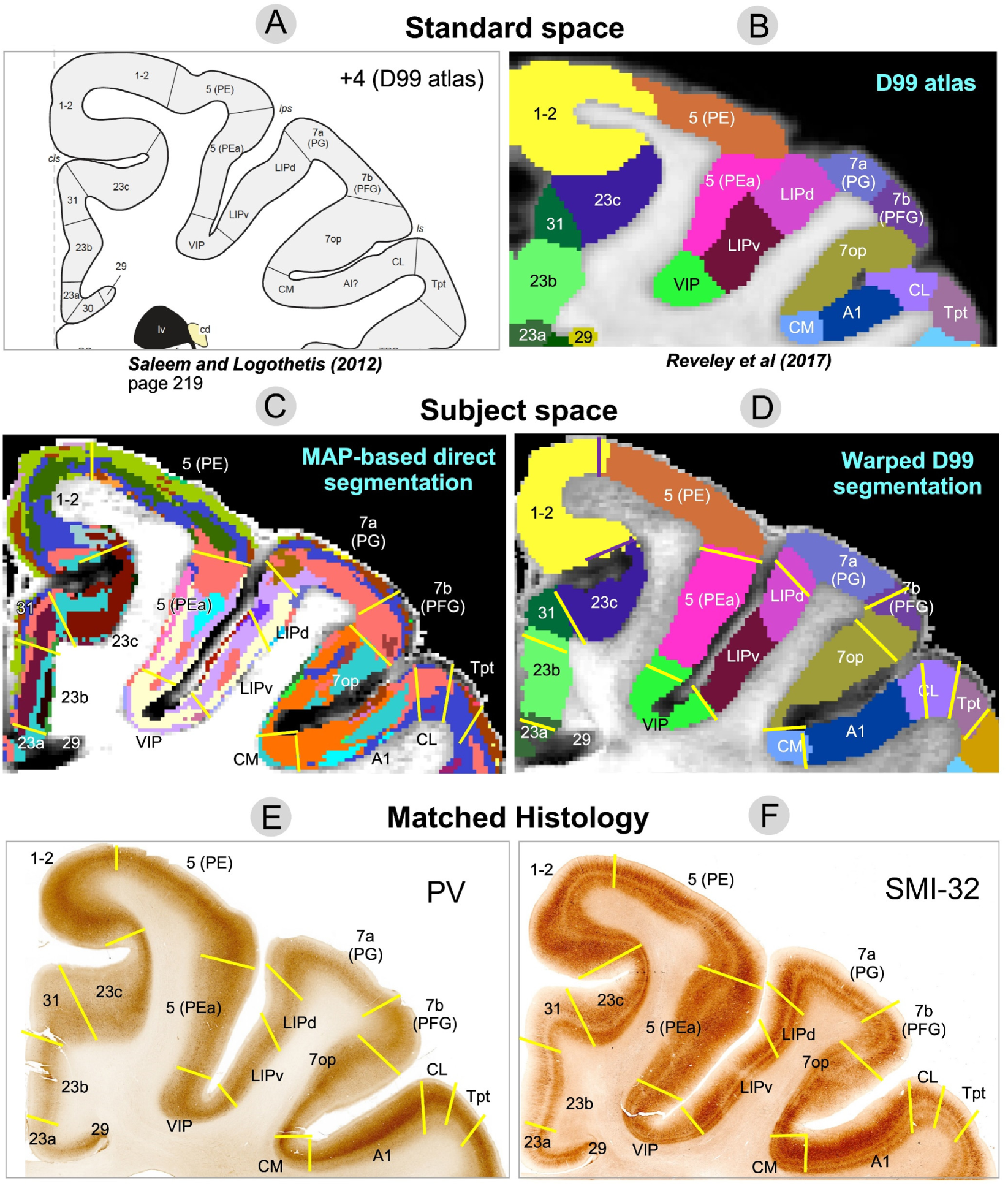
Comparison of warped D99 and the direct MAP-based segmentations with two histological stains in a matched brain region. Boundaries between cortical areas were delineated manually using only the histological images (E and F, yellow and black lines). Discontinuities in the laminar patterns of the MAP-based segmentation (C) correspond well with those observed in the PV (E) and SMI-32 (F) matched histological slices, providing a more accurate estimation of areal boundaries than the warping-based segmentation (D); e.g., the borders between areas 7op and 7b(PFG), LIPd and 7a(PG), A1 and CL, CL and Tpt, among others. In regions with high cortical curvature, all three methods showed good agreement; e.g., borders between areas 1-2 and 23c, 5(PE) and 5(PEa).

Although laminar patterns observed with MAP-MRI parameters may not consistently match those observed with any specific histological stain, the transition regions between laminar patterns observed with MAP-MRI and histology can be accurately identified and matched. Laminar pattern discontinuities identify borders between cortical areas and can be compared to similar estimates obtained with warping-based cortical parcellation and histology. In general, transition regions between MAP-derived laminar patterns can be matched well to the areal borders estimated with the warping-based D99 cortical parcellation, although the results of direct MAP-based clustering appear to be more consistent with the cytoarchitecture observed in the corresponding histological images (Fig. 7-8). This consistency could be traced across successive coronal sections of the stained tissue. Although the MRI-derived segmentations yielded volume information (i.e,. 3D), the 2D nature of histological analysis in our study restricts the comparison to coronal sections only.

### 3.5 MAP-based estimation of areal borders

Figures 7 and 8 illustrate two examples comparing the direct MAP-based segmentation and the warping-based D99 parcellation with matched histological coronal sections. Areal borders delineated manually based on discontinuities in the laminar patterns observed on the PV (Figs. 7E, 8E) and SMI-32 (Figs. 7F, 8F) stained sections (yellow lines) were superimposed on the two MRI-derived segmentations (Figs. 7C, 7D) and warping-based atlas parcellations (Figs. 7D, 8D). The transition regions between laminar patterns in the MAP-based segmentation (Fig. 7C) correspond well with those observed in histological sections, providing more accurate estimates of areal borders than the warping-based cortical parcellation.

For instance, the abrupt change in intensity in the PV-stained section (Fig. 7E) that defines the border between areas F1(4) and 3a/b matches well with the laminar pattern discontinuity obseerved on the MAP-based segmentation (Fig. 7C). Similarly, the intensity change in the SMI-32-stained section (Fig. 7F) corresponding to the border between areas VIP and 5(PEa) can be easily matched with the corresponding laminar pattern transition region in the MAP-based segmentation (Fig. 7C). In both cases, these areal borders located along sulcal walls (i.e., almost no curvature) are clearly more accurately delineated in the MAP-based segmentation (Fig. 7C) than in the warping-based D99 parcellation (Fig. 7D). In Fig. 8, the MAP segmentation (Fig. 8C) shows a laminar pattern discontinuity that correlates well with the signal intensity gradients in both PV-(Fig. 8E) and SMI-32-stained sections (Fig. 8F) delineating the border between areas 7op and 7b(PFG) (Fig. 8, yellow lines) more accurately than the warping-based D99 cortical parcellation (Fig. 8D). Similarly, the border between areas A1 and CL delineated by the signal intensity discontinuity on the SMI-32-stained section (Fig. 8F) can be more accurately traced on the MAP-based segmentation (Fig. 8C) than on the warping-based D99 cortical parcellation (Fig. 8D).

The D99 atlas labels from standard space (Figs. 7B, 8B) were warped to the subject space based on the deformation field produced by the nonlinear registration between the subject’s structural scan and the standardized template accompanying the atlas. This registration typically relies on structural scans with good GM/WM contrast, such as T1W, T2W, or MTR volumes. Consequently, the subject-specific anatomical landmarks that drive the nonlinear registration are not mesoscopic cytoarchitectonic features (e.g., cortical layers) but the GM/WM and GM/CSF boundaries, i.e., the WM and pial surfaces, respectively, which determine the 3D geometry of the cortical ribbon. Therefore, not surprisingly, the best correspondences between the warped D99 areal boundaries and the matched histology were generally found in regions with high cortical curvature. For instance, the borders between areas 1-2 and 3a/b, and between areas 5(PE) and 5(PEa), which follow gyral crests (high convexity) in the histological images (Fig. 7E-F) are accurately estimated by the warped D99 segmentation (Fig. 7D). We can also clearly observe these areal boundaries in the MAP-based segmentation as abrupt discontinuities in the corresponding laminar patterns (Fig. 7C), also in excellent agreement with histology. Similarly, in Fig. 8, the border between areas 1-2 and 23c, which follows a sulcal bottom (high concavity) in histology (Fig. 8E-F), is accurately delineated by both warped D99 (Fig. 8D) and direct MAP-based segmentation (Fig. 8C). On the other hand, we found the largest discrepancies between the MAP and D99 segmentations when comparing some areal borders located in regions with lower cortical curvature, such as the sulcal walls (e.g., the F1(4)-3a/b and VIP-5(PEa) borders in Fig. 7, or the 7op-7b(PFG) border in Fig. 8). As expected, the lack of subject-specific anatomical landmarks in regions with low cortical curvature (e.g., sulcal walls) reduces the accuracy of warping-based cortical parcellation with conventional structural scans. Conversely, direct MAP-based segmentation is sensitive to intrinsic cytoarchitectonic features and can identify these borders more accurately in single individuals.

To enable a quantitative estimation and comparison of distances between the areal borders estimated using the three methods, Fig. SA1 in Supplementary Material A shows the same regions from Figs. 7 and 8, respectively, but overlays areal borders estimated from MAP and histology on the warping-based segmentation, and includes a scale bar. This analysis provides a more direct and quantitative assessment of the segmentation accuracies but is still limited to 2D sections. Within a 2D section, the orientations of the estimated areal borders from the three methods can differ. Consequently, the relative distances between the corresponding borders must be measured at a consistent cortical depth, e.g., the mid-cortical layer. This inadequacy can reduce accuracy and degrade the significance of the quantitative assessment. The quantitative comparison of MAP-based and conventional warping-based cortical segmentations with the corresponding histology may be affected by several sources of error, including imperfect registration/matching between MRI and histology (Pas et al., 2020); variability in histological processing of whole-brain sections: tissue deformation, cutting, folding, and tearing during the preparation of microscope slides (i.e., slicing, staining, and mounting), spatial variations in stain concentration; and difficulty following structures/layers/domains in 3D across stained sections. In future studies, it may be possible to overcome many of these impediments using advanced methods for 3D registration between MRI volumes and histological sections (Wagstyl et al., 2020) and automated algorithms for reliably detecting laminar patterns and their discontinuities (Schiffer et al., 2021; Schleicher et al., 2005; Wagstyl et al., 2020).

In Supplementary Material B, we replicated Figs. 5-8 using GMM clustering based solely on DTI microstructural parameters (FA, RD, and AD). Although DTI-based GMM analysis resulted in a lower BIC than MAP-based analysis, the DTI-based segmentation showed more fragmented cortical layers (Fig. SB1A-B) and asymmetry between the left and right hemispheres (Fig. SB2), indicating inconsistencies in the size, shape, and location of cytoarchitectonic clusters. DTI parameters are generally less robust than MAP parameters in many cortical regions due to the limitations of the tensor model in capturing diffusion anisotropy, particularly in cortical areas with varying proportions of radial and tangential diffusion components(Avram, Saleem, Komlosh, et al., 2022). For example, the borders between areas 1-2 and 3a/b, areas F1(4) and 3a/b, or between VIP and AIP align more accurately with the laminar pattern discontinuities from the MAP-derived segmentation (Fig. 7C) than with those from the DTI-derived segmentation (Fig. SB3C). Nevertheless, the discrepancies between DTI (Figs. SB3 and SB4) and MAP-based parameters (Figs. 7 and 8) were relatively minor in our study (Supplementary Material B), likely due to the high spatial resolution (200 *µ*m), which reduces tissue heterogeneity within each voxel and reduces the discrepancies between the tensor model and the MAP signal representation (Avram, Saleem, & Basser, 2022).

## 4 Discussion

Recent advances in MRI technology (D. Feinberg et al., 2021; Huang et al., 2021) have continuously enhanced the spatial resolution of clinical structural scans, including MPRAGE, MTR, T_1_W, and T_2_W MRI, bringing it increasingly closer to the mesoscopic scale. However, despite these notable improvements in spatial resolution and signal-to-noise ratio (SNR) efficiency, these scans offer limited additional contrast for detailed cortical structures, such as layers and areal borders. As a result, their accuracy in warping-based cortical parcellation applications is only marginally improved. In this study, we show that high-resolution MAP-MRI parameters are particularly sensitive to intrinsic cytoarchitectonic features, allowing for the direct detection of boundaries between cortical domains with different microstructures and architectures, in good agreement with histology. These robust and reliable quantitative biomarkers can be acquired noninvasively with excellent reproducibility (Avram et al., 2016) and will likely play a unique and crucial role in the development of more precise and individualized methods for structural segmentation and characterization of the cortex. MAP-based cytoarchitectonic segmentation could complement and augment histological tissue characterization, refine borders between cortical areas estimated with conventional warping-based cortical parcellation, and enable the construction of a digital 3D cytoarchitectonic mesoscale brain atlas for use in neuroscience research and clinical applications (Amunts et al., 2013; Howard et al., 2023; Paquola et al., 2021; Wagstyl et al., 2018).

### 4.1 Denoising using the cortical reference frame

The local orientation of the cortical surface can be used to effectively denoise ultra-high-resolution images and improve visualization of signals from cortical layers (L. R. Huber et al., 2021; Wagstyl et al., 2018). In this study, we have applied this principle to denoise high-resolution cortical dMRI data by CRF-filtering each distortion-corrected and co-registered DWI individually, in a separate step that precedes the microstructural analysis, i.e., the estimation of diffusion propagators and MAP parameters. Unlike MAP parameter volumes, DWIs are generally sparse and can have sharp and pronounced edges in regions of anisotropy, especially at higher b-values. These edges are aligned with the orientations of anisotropic structures, such as cortical layers. By filtering each DWI separately using an anisotropic Gaussian filter whose orientation is informed by the local anatomy (i.e., CRF orientation), we can effectively reduce noise while preserving laminar contrast.

The CRF-filtering step can be applied to denoise any ultra-high-resolution cortical dMRI dataset and may be retrospectively applicable to data from other studies. It requires, at each voxel, the estimation of the parallel and perpendicular orientations to the cortical surface, i.e., the cortical reference frame. We employed a straightforward approach to estimate these orientations using LayNii directly from the MTR volume after co-registration with the dMRI data (Fig. 1). Alternatively, the CRF may be derived by estimating the pial and white matter surfaces with Freesurfer (Fischl, 2012) and interpolating the direction normal to the cortical surface at each voxel based on topologically consistent surface meshes sampling different cortical depths (Avram et al., 2020a, 2020b). Finally, the correspondence between the diffusion tensor reference frame and the cortical reference frame provides yet another method for estimating the radial and tangential cortical orientations in each voxel, in this case, directly from the dMRI data itself (Avram et al., 2020a, 2020b). Under normal conditions (i.e., healthy brain) and at sufficiently high spatial resolutions (Avram, Saleem, & Basser, 2022), these three methods are expected to yield consistent CRF estimates throughout most of the cortex in agreement with histological observations of cytoarchitecture (Avram, Saleem, & Basser, 2022; Avram et al., 2020a, 2020b).

### 4.2 MAP-MRI-based direct 3D cortical segmentation

The clustering method used in this study, Gaussian mixture modeling, is a straightforward and appropriate choice for our dataset. GMM is particularly effective for clustering large data sets with complex data distributions, as it can accommodate clusters of varying sizes and variances. Moreover, the convergence of the GMM algorithm is unaffected by the order of the input data, ensuring consistent cluster assignments even if voxel positions are randomly permuted. Specifically, at each iteration the expectation-maximization (EM) algorithm that optimizes the GMM likelihood processes the entire dataset as a batch, using calculations over the complete dataset (posterior probabilities and aggregated statistics) to update the model parameters. These operations are independent of the order of the data points, yielding the same results even if the sequence of data points is randomly shuffled. The convergence of the EM algorithm is determined based on the overall likelihood function which is also independent of the order of the data points, yielding the same convergence behavior and final parameter estimates regardless of the ordering of the input data points (McLachlan & Basford, 1988).

Direct voxelwise clustering of MAP parameters could be improved using more advanced methods such as spectral or hierarchical clustering or algorithms relying on machine learning and artificial intelligence. However, these methods generally require larger computational resources (CPU processors, memory, and speed) and/or separate training steps using validated data. It is important to highlight that the GMM clustering analysis used exclusively scalar-valued intrinsic MAP/DTI microstructural parameters with no information about voxel positions.

The 3D morphological filtering step uses the relative positions of clustered voxels to uniquely relabel spatially disjoint components (groups of voxels) that were initially assigned to the same cluster and to remove or merge small isolated components into larger neighboring clusters. This process increases the total number of distinct labels and may result in some labels having similar MAP biomarker profiles. However, the MAP profiles of each label remain distinct from those of their neighboring clusters. Therefore, this step improves the visualization of the cytoarchitectonic borders between domains with different MAP biomarker fingerprints. Additionally, 3D morphological processing reduces the sensitivity of the GMM clustering analysis to the initial number of clusters, providing a relatively consistent number of unique segmented cytoarchitectonic domains in the final output.

After GMM clustering and 3D morphological filtering, the labels of the segmented cytoarchitectonic domains were arbitrarily assigned. The best matching of labels across the left and right hemispheres was achieved by computing the cross-tabulations of the MAP segmentation and the symmetric D99 atlas in each hemisphere and then solving a minimum cost perfect matching problem efficiently and exactly using the Kuhn-Munkres assignment algorithm (Kuhn, 1955; Munkres, 1957). In this final step, the shapes and boundaries of the segmented domains do not change. Instead, information from the symmetric D99 atlas is used to assign consistent labels (colors) to the corresponding regions in the left and right hemispheres (Fig. 2).

The manual method of defining the histological areal boundaries in our analysis is prone to errors due to tissue deformations during histological processing and due to the imperfect matching between the 2D sections and the 3D MRI volumes. One limitation of our current study is the lack of 3D co-registration between MRI and histology. The use of multiple stains allows more reliable delineation and validation of many areal boundaries across the entire cortex but makes 3D registration and analysis of histological data even more challenging. Unlike other studies (Wagstyl et al., 2020), the histological analysis in this study was not specifically designed for 3D whole-brain registration.

### 4.3 Improving the MAP-based cytoarchitectonic segmentation

Our results are in good agreement with findings from other studies that have explored the potential of high-resolution dMRI as a noninvasive contrast for cortical segmentation (Bastiani et al., 2016; Little & Beaulieu, 2021; Nagy et al., 2013; Oros-Peusquens et al., 2012), etc. While previous studies focused on voxel segmentation in specific parts of the cortex and/or acquired dMRI data (e.g., single-shell data) suitable for a specific dMRI analysis (e.g., DTI, HARDI), in our study, we directly cluster voxels across the entire cortex based on quantitative features of their diffusion propagators. For example, studies have shown that k-means clustering applied to high angular diffusion MRI (HARDI) data can identify borders between cortical areas (Bastiani et al., 2016; Nagy et al., 2013) and distinguish between superficial and deep layers (J. Zhang et al., 2022). In these studies, the voxels were classified according to their orientational diffusion characteristics (i.e., FODs) as determined by the corresponding spherical harmonic coefficients. However, these features, when defined in the laboratory reference frame, can vary with the local curvature of the cortical ribbon, potentially biasing the results (Bastiani et al., 2016; Cottaar et al., 2018; Kleinnijenhuis et al., 2015; Nie et al., 2011). In our study, we use multishell dMRI data to explicitly and reliably measure diffusion propagators using MAP-MRI. We demonstrated for the first time that cortical domains with distinct cytoarchitectonic features can be delineated by directly clustering scalar microstructural parameters that quantify *intrinsic* salient propagator features effectively and comprehensively in the local cortical reference frame of each voxel. Moving forward, the MAP-based cytoarchitectonic segmentation may be improved by:

1. using more advanced clustering algorithms
2. incorporating orientational features (e.g., DEC-FOD image)
3. integrating structural connectivity information for each voxel, i.e., cortico-cortical connectivity (Anwander et al., 2007; Barbas & Rempel-Clower, 1997; Fan et al., 2016; Gao et al., 2018; Johansen-Berg & Rushworth, 2009; Klein et al., 2007; Leuze et al., 2014; Moreno-Dominguez et al., 2014). In principle, this could be computed from the same high-resolution MAP-MRI data set using fiber tractography. However, unlike simple GMM clustering of voxelwise values, the connectivity between cortical voxels (areas and layers) introduces non-local information that may be biased by cumulative errors during fiber tracking.
4. incorporating functional cortico-cortical connectivity information. Layer-specific fMRI could provide additional features for identifying distinct cortical substructures based on their function and connectivity (L. Huber et al., 2021; Shipp, 2007).
5. integrating spatial information, e.g., the spatial location of each voxel or a reference topology of cortical layers and areas
6. averaging multiple datasets to construct a standardized template of diffusion propagators and MAP parameters at the mesoscopic scale (Avram, Bernstein, Irfanoglu, et al., 2018; Avram, Bernstein, Irfanoglu, et al., 2019). Despite its indisputable scientific value, such a project is difficult to undertake because of the long scan durations and attendant costs.
7. incorporating additional salient quantitative imaging parameters or biomarkers and correlations between them, such as T_1_, T_2_, MTR, spectral relaxation information (Avram, Magdoom, et al., 2022; Avram, Saleem, & Basser, 2022; Avram, Sarlls, & Basser, 2019, 2021; Benjamini & Basser, 2020), or morphological features in the clustering analysis.

### 4.4 The role of MAP-MRI in cortical segmentation

Table 1 compares direct MAP-based segmentation with conventional warping-based methods and histological tissue analysis. The biophysical contrast mechanisms of these approaches characterize the cortical anatomy at different length scales. Histological analysis elucidates cellular and subcellular structures at the microscopic scale, conventional structural MRI outlines the 3D shape of the entire cortical gray matter at the macroscopic scale, and MAP-MRI provides intrinsic cytoarchitectonic contrast at the mesoscopic scale, bridging the gap between the other two methods. Therefore, MAP-based segmentation can improve both conventional warping parcellation and histological tissue analysis, while providing a 3D noninvasive measurement.

**Table 1:**
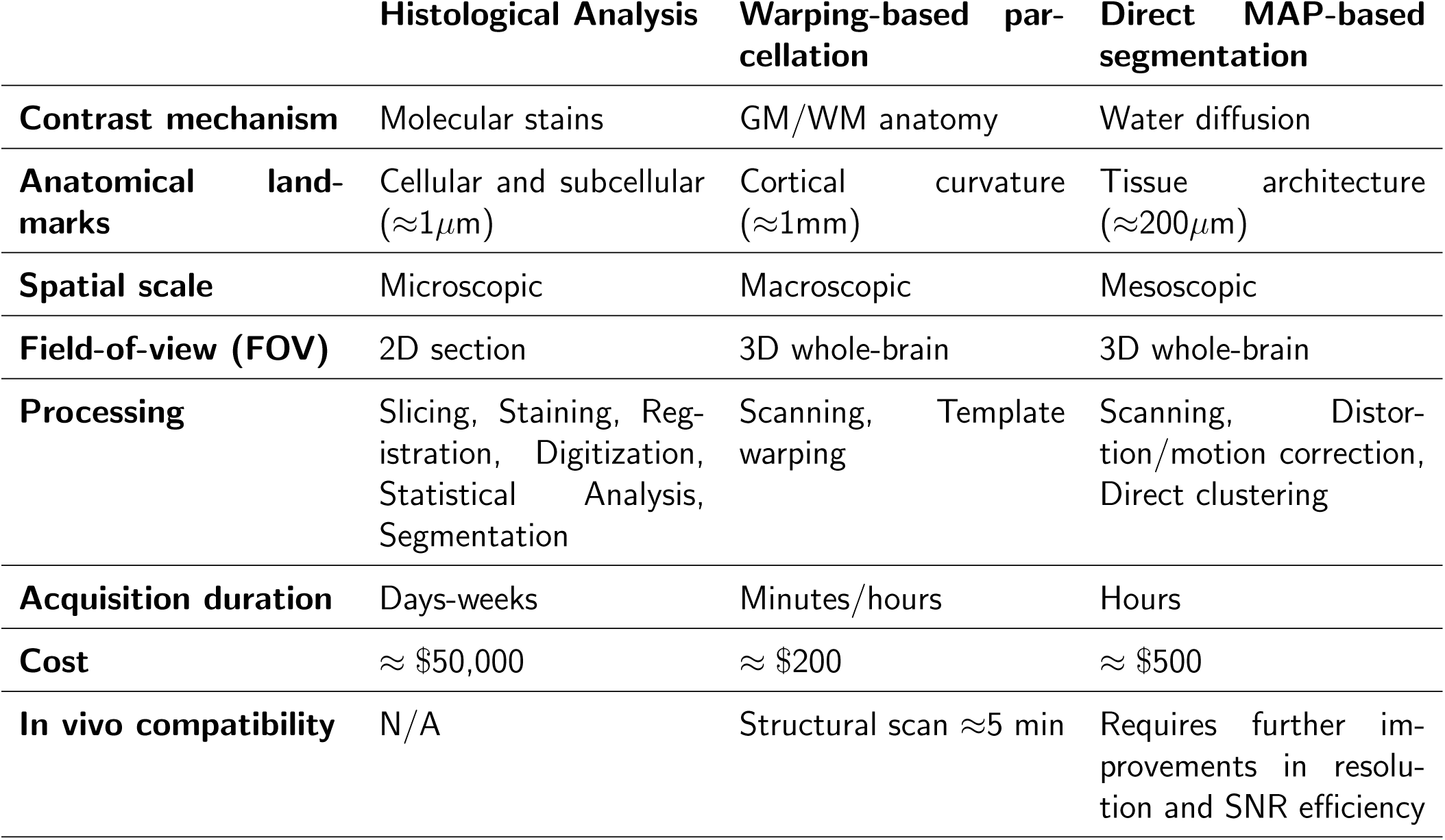
Advantages and disadvantages of three approaches for whole-brain cortical segmentation/parcellation. Research activities focus on improving the spatial resolution and SNR efficiency of *in vivo* high-resolution dMRI acquisitions (D. A. Feinberg et al., 2023; Huang et al., 2021; F. Wang et al., 2021), including MAP-MRI (Huang et al., 2021), and on developing automated processing pipelines and computational methods for 2D and 3D analysis of histological data (Schiffer et al., 2021; Schleicher et al., 2005; Wagstyl et al., 2020).

Using state-of-the-art histology, microscopy, and machine learning techniques, it is possible to significantly improve the detail and accuracy of cortical atlases. Wagstyl et al. (Wagstyl et al., 2020) for instance used convolutional neural networks to automatically segment cortical layers from contiguous histological sections in an entire human brain (Amunts et al., 2013). Such detailed datasets cannot be acquired in individual live subjects, and the painstaking and costly analysis of whole brain histological data can be prohibitive for studies using a large number of brain specimens. However, such studies produce extraordinary reference datasets that can be used to construct 3D histological human cortical atlases that enable large-scale indirect cortical layer segmentation based on warping the template to easily-acquired T_1_-weighted MRI scans of single individuals (Paquola et al., 2021). Although the resolution of these atlases is exquisite, their accuracy in subject-specific applications can be blunted by the lack of cortical cytoarchitectonic contrast in both the subject’s MRI scans and the standardized template.

During warping-based parcellation anatomical features at different spatial scales in the subject and template MRI volumes are aligned. Since even at ultrahigh spatial resolution these volumes (T1-weighted or T2-weighted) do not reveal intrinsic cortical features (e.g., laminar patterns and areal borders) the registration of cortical voxels is driven primarily by extrinsic cortical features, such as the high-contrast interface between cortical GM and CSF and cortical GM and WM, i.e., the pial and WM surfaces. To overcome this limitation, high-resolution MRI techniques that provide new contrasts sensitive to cortical microstructural features, like MAP-MRI, are needed. The inherent microstructural sensitivity of MAP biomarkers provides an alternative means for developing ultrahigh (mesoscopic) resolution 3D cytoarchitectonic atlases (Saleem et al., 2021, 2023, 2024) and templates to enable more accurate propagator-based warping (Avram, Bernstein, Irfanoglu, et al., 2019) of similar MAP-MRI data sets from single individuals based on both extrinsic and intrinsic cortical anatomical features.

The MAP parameters (e.g., PA, NG, RTOP) derived from the normalized dMRI signals are less sensitive to proton density, B_1_, or B_0_ inhomogeneities. Compared to other MRI contrasts, MAP parameters have reduced sensitivity to experimental factors such as TE or TR and high reproducibility *in vivo* (Avram et al., 2016). They reliably and robustly quantify the salient characteristics of the diffusion propagator that describe the average diffusive motion of all water molecules in each voxel. Unlike histological stains, which label specific proteins, MAP parameters quantify important differences in the voxel-averaged tissue microstructure and cytoarchitecture, capturing information about the densities, sizes, and shapes of cells, their distribution patterns, and dominant orientations of their arrangements, their membrane permeabilities, and the densities of neurofilaments and other subcellular structures that may impede water diffusion. Consequently, two cytoarchitectonic domains with very similar MAP fingerprints (i.e., MAP-derived parameter values) might have significantly different intensity patterns than PV or SMI-32 stains (Avram, Saleem, Komlosh, et al., 2022). Nevertheless, our findings suggest that spatial changes in MAP-derived laminar patterns correspond well with transition regions in cortical cytoarchitecture observed with histology, making high-resolution MAP-MRI suitable for cytoarchotectonic parcellation, despite its lack of molecular specificity.

Despite its cellular and subcelluar spatial resolution, histological analysis of large brain regions produces extensive data sets and often requires substantial computational resources for automated processing, including 3D registration, downsampling, and statistical analysis. Advances in automated processing pipelines and computational methods for 2D and 3D histological data analysis (Schiffer et al., 2021; Schleicher et al., 2005; Wagstyl et al., 2020) have significantly enhanced our ability to manage large datasets and extract essential features for automatic cortical segmentation. On the other hand, MAP-derived parameters inherently average microstructural characteristics over mesoscopic length scales (*≈*200*µm* voxel), providing manageable datasets that are more suitable for distinguishing cytoarchitectonic domains across the entire cortex. The microstructural specificity of MAP parameters complements the molecular specificity of histological staining for tissue analysis and provides a convenient and efficient method for cytoarchitectonic mapping. This study illustrates the ability of diffusion MRI to bridge multiple length scales, ranging from microscopic diffusion distances *≈*1-7*µm* experienced by individual water molecules, to mesoscopic cytoarchitectonic domains of *≈* 100-700*µm*, to macroscopic dimensions of imaging voxels and the cortex as a whole *≈*1-7cm.

### 4.5 Potential for clinical translation

The mesoscopic resolution required for direct MAP-based cytoarchitectonic cortical segmentation is on the order of the thickness of a cortical layer, approximately 400*µm*. Using state-of-the-art technology, whole-brain acquisitions at such ultrahigh spatial resolutions have been achieved in live human subjects using T1-, T2-, T2*-weighted contrasts (D. A. Feinberg et al., 2023; Huang et al., 2021), but not yet for dMRI. At this resolution, dMRI requires significantly higher SNR levels and longer scan durations. Moreover, the efficient 3D acquisition of volumes with a large imaging matrix (i.e., whole-brain and high-resolution) using segmented k-space trajectories is nicely compatible with fMRI, but not with *in vivo* dMRI, or MAP-MRI. *In vivo* dMRI with 3D acquisition requires cardiac gating, the use of navigation, and/or other specialized acquisition and postprocessing methods (Avram et al., 2014) to correct for shot-to-shot random phase variations induced by tissue motion. Although whole-brain *in vivo*dMRI acquisitions are currently limited to *≈* 800*µm* resolution and prohibitively long scan durations (F. Wang et al., 2021), localized dMRI acquisitions with the mesoscopic resolution required for resolving individual cortical layers may be achievable in clinical scans with a reduced FOV and reasonable scan durations (Balasubramanian et al., 2021; Heidemann et al., 2012; Holdsworth et al., 2019).

Collaborations between research groups and scanner manufacturers aim to improve the resolution for *in vivo* dMRI using various avenues, including developing high-field MRI systems (D. Feinberg et al., 2021), high-performance gradients (D. Feinberg et al., 2021; Huang et al., 2021; McNab, Edlow, et al., 2013), innovative RF technology (Keil et al., 2013; Truong et al., 2014), clever strategies for more accurate and accelerated dMRI acquisition (Avram et al., 2014; Balasubramanian et al., 2021; Dong et al., 2024; D. A. Feinberg et al., 2010; Setsompop et al., 2012; F. Wang et al., 2019), more efficient dMRI sampling schemes (Afzali et al., 2021; Avram, Sarlls, et al., 2018), and improved pre– and post-processing pipelines to remedy various imaging artifacts (Tax et al., 2022). In parallel, surface-based analysis (Avram, Saleem, & Basser, 2022; McNab, Polimeni, et al., 2013) and multidimensional diffusion encoding (Topgaard, 2017; Westin et al., 2016) may allow us to quantify microscopic cortical components in healthy individuals (Avram, Sarlls, & Basser, 2019, 2021; Avram et al., 2010; Magdoom et al., 2023) and patient (Lampinen et al., 2020) populations with standard spatial resolution that can be acquired on clinical scanners (Avram, Magdoom, et al., 2022; Avram, Tian, et al., 2021). These developments support the creation of MAP-based whole-brain cytoarchitectonic templates and atlases (Saleem et al., 2021, 2023, 2024) for use in individual subjects as well as group/longitudinal/cross-sectional studies (Avram, Bernstein, Irfanoglu, et al., 2019).

Our study highlights the use of high-resolution MAP-MRI as a noninvasive, sensitive tool with intrinsic cytoarchitectonic contrast for subject-specific cortical segmentation, marking a significant step toward developing MAP-based whole-brain cytoarchitectonic templates and atlases (Saleem et al., 2021, 2023, 2024). Identifying cortical laminar patterns enhances structural cortico-cortical connectivity mapping (Leuze et al., 2014) and could provide valuable functional insights for neuroscience and clinical applications (Avram & Basser, 2014; Shipp, 2007; Weiler et al., 2008). Integrating MAP biomarkers with histological analysis has potential applications in the study of aging (Bouhrara et al., 2023), traumatic brain injury (Hutchinson et al., 2018), epilepsy (Chen et al., 2019), cancer (She et al., 2023; P. Wang et al., 2022), multiple sclerosis (Brusini et al., 2021), stroke (Boscolo Galazzo et al., 2018), Alzheimer’s disease (Spotorno et al., 2022), Parkinson’s disease (Le et al., 2020), and other neurological conditions.

## 5 Conclusions

Our results demonstrate that direct clustering can effectively distinguish 3D cortical domains at the mesoscale based on cytoarchitectonic differences as measured by MAP parameters. We observed laminar patterns whose discontinuities helped delineate subject-specific borders between cortical areas more accurately than conventional warping-based methods when compared with the corresponding histological sections from the same brain. The compelling sensitivity of high-resolution MAP-MRI parameters to intrinsic cytoarchitectonic features (Avram, Saleem, Komlosh, et al., 2022) could complement and augment existing methods for tissue characterization, such as histological analysis and warping-based cortical parcellation, to enable an accurate “personalized” (i.e., subject-specific) cortical segmentation.

## Data and Code Availability

The MATLAB routines used in this project are publicly available at https://github.com/4lexandru/ MAPMRI segmentation. The TORTOISE software package used for registration and distortion correction of diffusion MRIs is freely available at https://tortoise.nibib.nih.gov/. The ANTs software package is freely available at https://stnava.github.io/ANTs/. The FSL software package used for brain tissue segmentation is freely available at https://fsl.fmrib.ox.ac.uk/fsl/fslwiki/FslInstallation. The LayNii software package used for the estimation of cortical layers is freely available at https://github.com/layerfMRI/ LAYNII.

## Author Contributions

**Kristofor E. Pas**: Data curation, Formal analysis, Investigation, Methodology, Software, Visualization, Writing – review & editing;

**Kadharbatcha S. Saleem**: Data curation, Formal analysis, Investigation, Methodology, Resources, Validation, Visualization, Writing – review & editing;

**Peter J. Basser**: Funding acquisition, Investigation, Methodology, Project administration, Resources, Writing – review & editing;

**Alexandru V. Avram**: Conceptualization, Data curation, Formal analysis, Investigation, Methodology, Project administration, Software, Supervision, Visualization, Writing – original draft, Writing – review & editing.

## Acknowledgements

We thank Drs. Michal Komlosh, Cecyl Chern-Chyi Yen, and Frank Ye for assistance with sample preparation and data acquisition, and Drs. Bernard Dardzinski and Alexandru Korotcov for providing the RF coil used in this experiment. We thank Tom Pohida and Marcial Garmendia-Cedillos for their help in constructing the 3D-printed brain mold. We also thank Drs. Ted Usdin and Sarah Williams-Avram as well as Drs. Vincent Schram and Ross Lake for help with optical imaging and digitization of microscope slides, and FD Neurotech for histological services provided. We also thank Drs. Betsy Murray and Richard Sanders from the Laboratory of Neurophysiology at NIMH for providing the perfusion-fixed monkey brains for our experiments. This work utilized computational resources of the NIH HPC Biowulf cluster (http://hpc.nih.gov).

The opinions expressed herein are those of the authors are not necessarily representative of those of the Uniformed Services University of the Health Sciences (USUHS), the Department of Defense (DoD), VA, NIH, or any other US government agency, or the Henry M. Jackson Foundation.

## Supplementary Material

Supplementary Material (created during production as a web link to online material).

